# High-throughput cryo-EM structure determination of amyloids

**DOI:** 10.1101/2022.02.07.479378

**Authors:** Sofia Lövestam, Sjors H.W. Scheres

## Abstract

The formation of amyloid filaments is characteristic of various degenerative diseases. Recent breakthroughs in electron cryo-microscopy (cryo-EM) have led to atomic structure determination of multiple amyloid filaments, both of filaments assembled *in vitro* from recombinant proteins, and of filaments extracted from diseased tissue. These observations revealed that a single protein may adopt multiple different amyloid folds, and that *in vitro* assembly does not necessarily lead to the same filaments as those observed in disease. In order to develop relevant model systems for disease, and ultimately to better understand the molecular mechanisms of disease, it will be important to determine which factors determine the formation of distinct amyloid folds. High-throughput cryo-EM, in which structure determination becomes a tool rather than a project in itself, will facilitate the screening of large numbers of *in vitro* assembly conditions. To this end, we describe a new filament picking algorithm based on the Topaz approach, and we outline image processing strategies in Relion that enable atomic structure determination of amyloids within days.

## Introduction

Amyloids are filamentous, helical aggregates of proteins that are characterised by a cross-β-sheet quaternary structure. Amyloid formation of a few dozen proteins in the human genome is associated with more than 50 diseases (Knowles, Vendruscolo and Dobson, 2014)

Historically, structure determination of amyloids has been more difficult than for globular proteins. Many globular proteins can form crystals, making them amenable for X-ray diffraction studies. Alternatively, solution-state nuclear magnetic resonance (NMR) can be used for relatively small proteins. However, often the helical symmetry of amyloids is incompatible with crystallisation, and their size precludes solution-state NMR. Before the advent of atomic structure determination by cryo-EM, amyloids were mainly studied by solid-state NMR, which requires large amounts of ^13^C ^15^N labelled protein (Tycko, 2006; Tuttle *et al.*, 2016).

Atomic structure determination of globular proteins by cryo-EM became mainstream through direct electron detectors and statistical image processing software (Bai *et al.*, 2013; Li *et al.*, 2013). Application of the same technique to amyloids wasn’t successful until the implementation of helical symmetry in the Relion program (He and Scheres, 2017). Using this approach, the first cryo-EM structure of an amyloid was reported for filaments of the protein tau that were extracted from the brain of an individual with Alzheimer’s disease (Fitzpatrick *et al.*, 2017).

Besides Alzheimer’s disease, tau amyloid formation defines more than a dozen neurodegenerative diseases, which are collectively called tauopathies. Cryo-EM structures of tau filaments extracted from the brains of individuals with 13 different tauopathies revealed 8 different tau folds, and showed that different folds characterise different tauopathies (Falcon *et al.*, 2018a/b, 2019; Zhang *et al.*, 2020; Shi *et al.*, 2021). Multiple individuals with the same tauopathy showed the same folds. The factors that determine the remarkable structural specificity of tau folds among the different diseases remain unknown. Post-translational modifications, isoform composition, protein truncations, and protein or other co-factors may all play a role (Scheres *et al.*, 2020; Wesseling *et al.*, 2020; Limorenko and Lashuel, 2021).

Laboratory-based models, amenable to experimental perturbation, will be crucial to further our understanding of the molecular mechanisms of amyloid formation, which factors drive the formation of the different folds, and which role they play in disease. Full-length recombinant tau is extremely soluble, but it readily forms amyloids upon the addition of anionic cofactors such as heparin (Goedert *et al.*, 1996). Cryo-EM structures of heparin-induced recombinant tau showed that the resulting filaments are different from those observed in disease (Zhang *et al.*, 2019). Similar observations have also been made for other proteins. Filaments of amyloid-β (Kollmer *et al.*, 2019; Yang *et al.*, 2022) α-synuclein (Schweighauser *et al.*, 2020), serum amyloid A (Liberta *et al.*, 2019; Radamaker *et al.*, 2021) protein, TAR DNA-binding protein 43 (TDP-43) (Arseni *et al.*, 2021) and the prion protein (Kraus *et al.*, 2021; Manka *et al.*, 2021) that were extracted from diseased tissue were different from those assembled *in vitro*.

Recently, *in vitro* assembly conditions to replicate tau filaments of Alzheimer’s disease and chronic traumatic encephalopathy (CTE) were identified (Lövestam *et al.*, 2021). This effort involved solving 76 cryo-EM structures of recombinant tau filaments, including 29 structures that had not been observed previously. Different construct lengths, buffer conditions, shaking speeds and mutations mimicking post-translational modifications all affected the amyloid folds. Many of these folds could not be distinguished from the filaments observed in disease by negative stain EM or by cryo-EM two-dimensional (2D) class averaging.

We thus envision that large numbers of three-dimensional (3D) cryo-EM reconstructions will be required to further our understanding of the factors that drive the formation of specific amyloid folds, for tau as well as for other proteins. High-throughput methods for cryo-EM structure determination of amyloids will enable such studies. The first cryo-EM structures of tau filaments from the brain of an individual with Alzheimer’s disease (Fitzpatrick *et al.*, 2017) took over a year. Although the time required for amyloid structure determination has reduced since 2017, manual picking of filaments in the micrographs remained an important bottle neck in our recent work (Shi *et al.*, 2021; Yang *et al.*, 2022). Several automated approaches for helical filament picking have been proposed (He and Scheres, 2017; Huber, Kuhm and Sachse, 2018; Wagner *et al.*, 2019), but we find that manual picking often leads to better structures.

In this paper, we introduce a new method for automated picking of helical filaments. Our approach is a modification of the Topaz program, which employs a positive unlabelled deep-learning method that requires only few sparsely labelled particles for training and no labelled negatives (Bepler *et al.*, 2019). The Topaz program has been used successfully for cryo-EM structure determination of globular proteins, and it has recently been incorporated (through a wrapper) in Relion-4.0 (Kimanius *et al.*, 2021). We used the modified Topaz approach in our most recent work to automatically pick filaments for 38 out of the 76 *in vitro* assembled tau structures (Lövestam *et al.*, 2021).

Another bottle neck in cryo-EM structure determination of amyloids is initial model generation. Because of multiple local minima in the energy landscape of helical refinement, suboptimal initial models can lead to incorrect structures. We previously introduced a method that reconstructs initial 3D models from assembled 2D class averages and outlined the pitfalls of getting stuck in local minima during refinement (Scheres, 2020). Still, various users of our software have reported difficulties in solving amyloid structures. To facilitate high-throughput amyloid structure determination in more labs, we outline general processing strategies and highlight examples from three representative data sets of our recent work on recombinant tau (Lövestam *et al.*, 2021).

## Approach

### Automated filament picking in Topaz

The Topaz pipeline is composed of three steps (Bepler *et al.*, 2019). First, the micrographs are preprocessed. Relion’s wrapper performs a down-sampling and a normalisation operation that is equivalent to providing the `--affine` argument in Topaz. Second, a convolutional neural network is trained based on a small number of labelled particles and many unlabelled micrograph regions. Third, a sliding window is passed over the micrographs for classification with the trained network, and particle coordinates are extracted by non-maximum suppression. Our approach modifies only the coordinate extraction part of the third step and leaves the first two steps and the sliding-window part of the third step intact. The modified version outputs start-end coordinate pairs of straight segments of the filaments, which define the lines along which individual particle images are extracted and provide information for their orientational priors in Relion.

The sliding-window operation outputs an image with predicted scores for each position in the down-sampled micrograph. Our modification consists of four steps that operate on this image. First, the score image is binarized at a user-specified threshold (through the existing `-t` argument in Topaz). Second, the binarized image is skeletonised, using the `skeletonize` function (Zhang and Suen, 1984) from the scikit-image python package (Walt *et al.*, 2014). Third, straight lines are detected in the skeletonised image using the `probabilistic_hough_line` function (Galamhos, Matas and Kittler, 1999), again from scikit-image. This function takes three parameters: a threshold, a line length and a line gap. The line length can be controlled by the user through the newly implemented Topaz argument `--fl`. By default, the line length is set to twice the user-provided Topaz radius (which is set through the existing `-r` argument in Topaz). The threshold and the line gap parameters are fixed at 0.1 times the line length and the user-provided radius, respectively. Fourth, a custom-built algorithm merges lines into longer ones. Two lines are merged if the angle between them is smaller than 10° and two of their start or end coordinates lie within a distance from the other line that is smaller than the user-provided radius. The resulting start-end coordinate pairs can be used directly in Relion’s ‘Particle extraction’ jobtype.

Within Relion-4.0, training of the Topaz convolutional neural network and picking micrographs with the trained network are both performed with the ‘Auto-picking’ jobtype on the main graphical user interface (GUI). The Topaz radius (`-r`) is calculated as half the ‘Particle diameter’ that is specified on the Topaz tab of the GUI, taking the down-sampled pixel size into account. This diameter should reflect the average width of the filaments to be picked. When providing coordinates for the neural network training, it is important to provide coordinates for segments along the entire filaments, not only start-end coordinates. We recommend to manually pick start-end coordinates in a subset of the micrographs and to extract segments along these filaments using the ‘Particle extraction’ jobtype. The extracted `particles.star` file can then be used as the set of particles for Topaz training. If manual picking is not done precisely, or if individual segments along the lines defined by the start-end coordinates are not accurate because of curves in the filaments, a preliminary alignment of the extracted particles through the ‘2D classification’ jobtype may be used. In that case, the resulting `data.star` file is used as the set of particles for Topaz training. It is then important to check that the filaments are centred in the Y-direction of the 2D class averages.

The trained network may be used to auto-pick the rest of the micrographs. Filament picking is invoked through the `--f` argument. The most important parameter to tune is the binarization threshold (`--t`), with useful values in our work on recombinant tau ranging from −4 to −7. Tuning of the line length parameter of the Hough transform (`--fl`) is typically not necessary. These arguments are specified on the Relion-4.0 GUI in the ‘Additional Topaz arguments’ entry of the Topaz tab of the ‘Auto-picking’ jobtype. The resulting coordinates may be displayed using the ‘Display’ pull-down menu. In difficult cases, the argument `--fp` may be provided to launch a window that visualises the output of the four separate steps (binarization, skeletonization, Hough transform and line merging) for each micrograph (**Figure 1**). Based on these images, the user may then decide to tune the `-t`, `-r` or `--fl` arguments. If parameter tuning does not give satisfactory results, the network may need to be retrained.

**Figure 1:**
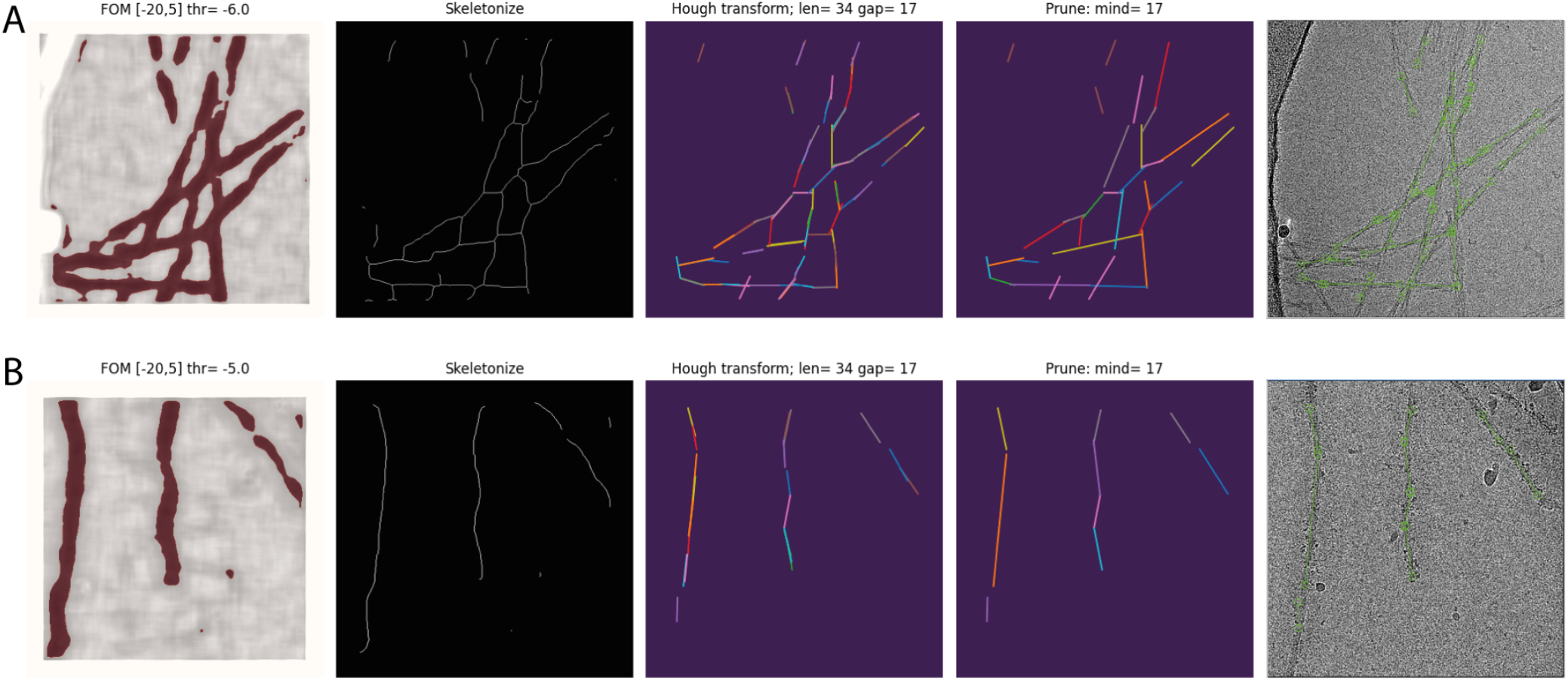
Automated filament picking. Rows A and B show the results for two different micrographs. The four panels from the left show the visualisation windows from the modified Topaz approach for the binarization, skeletonization, Hough transform and line merging steps, respectively. The panels on the right show the final start-end coordinates in the original micrograph.

The modified Topaz code is distributed under Topaz’s original GNU General Public Licence 3.0, as a fork of the original code at https://github.com/3dem/topaz.

### General strategy of amyloid structure determination in Relion

Structure determination of amyloids in Relion follows a broadly similar workflow as in conventional single-particle analysis of globular proteins. This section provides an overview of the general strategy and relevant differences between the two modalities. However, every data set is different, and it is hard to provide one approach that is suitable for all. Therefore, in the Results section we also describe in more detail relevant aspects of the processing of three of the data sets from our recent work on recombinant tau (Lövestam *et al.*, 2021). The first data set represents an easy case. Its structure was solved within 2 hours, while the data were still being acquired. The second data was more difficult due to the presence of different filament types with misleading crossover distances. The third data was the hardest, as it showed various filament types with almost identical 2D slices, encumbering their identification and separation. For each of these data sets, we share the original micrograph movies, together with all relevant intermediate results from their processing through the EMPIAR data base (Iudin *et al.*, 2016) (entries EMPIAR-10940, EMPIAR-10943 and EMPIAR-10944 respectively).

#### Micrograph inspection

Some data sets are not worth acquiring. Compared to many globular protein complexes, amyloids are relatively sturdy objects, but sometimes they do get damaged during sample preparation. Processing images of such filaments is likely to fail. One feature to look out for when acquiring data are what we call ‘swollen filaments’, which appear “blobby” with inconsistent widths along the filament axis, and result in poor 2D class averages. In addition, although it is in principle possible to solve the structure of filaments that do not twist, in practice this is often hampered by strong preferred orientations. **Figure 2** shows an example of good filaments, swollen filaments and non-twisting, flat ribbons.

**Figure 2:**
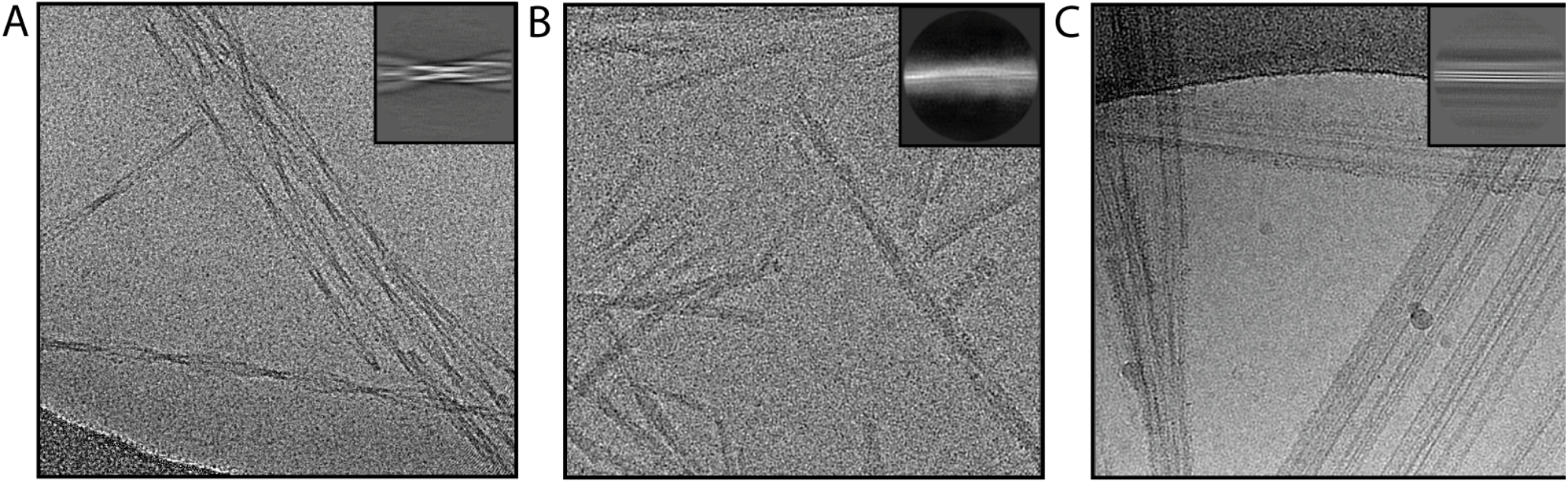
Representative micrographs. An example of micrographs is shown for good filaments (A), swollen filaments (B) and non-twisting ribbons (C). Insets on the top right show representative 2D class averages for the filaments shown.

#### Micrograph pre-processing and filament picking

Once suitable images have been recorded, micrograph movies are motion-corrected. We recommend using Relion’s own implementation of the UCSF MotionCor2 program (Zheng *et al.*, 2017; Zivanov *et al.*, 2018), which communicates metadata with particle polishing. Next, contrast transfer function (CTF) parameters are estimated, preferably using the open-source CTFFIND4 program (Rohou and Grigorieff, 2015). Filaments are picked manually in the micrographs by clicking start-end coordinate pairs or picked automatically using the modified Topaz approach described above. The start-end coordinate pairs define straight lines. Curved filaments are picked as multiple shorter lines. Individual particles images are extracted along these lines with an inter-particle distance that is defined by the helical rise (at this point one can provide a value of 4.75 Å) and the number of unique asymmetrical units (a value of 3 works well in most cases). It is often better to pick fewer, good filaments than larger numbers of suboptimal filaments. Filaments with anisotropically shaped cross-sections display alternatingly strong and weak signals in the micrographs. To avoid missing the weaker parts, relatively low Topaz threshold may be necessary. We recommend to visually check auto-picking results in several micrographs.

#### 2D classification

2D class averaging serves to assess the quality of the data, to remove suboptimal particles, and to detect the presence of multiple filament types. The latter requires user expertise if the filament types are similar. To aid novel users, we elaborate on this step for the second and third data sets that are described in the Results section. The mask diameter for 2D classification is set close to the box size; the angular sampling rate to 1-2°. The VDAM algorithm in Relion-4.0 (Kimanius *et al.*, 2021) works less well for amyloids as it does for globular proteins and better results are often obtained with the (slower) default algorithm.

It is often useful to calculate 2D class averages with a few different box sizes. Many final reconstructions are calculated in box sizes of approximately 250-300 Å, but earlier 2D classifications with box sizes in the range of 500-1000 Å are also useful. Although typically of lower resolution, larger 2D class averages facilitate identification of distinct filament types, can help measuring cross-over distances, and reduce the number of variables to optimise in the `relion_helix_inimodel2d` program for initial model generation.

#### Initial model generation

Subsets of 2D class averages that correspond to a single filament type are used to calculate initial 3D models in the `relion_helix_inimodel2d` program. This approach has been described in detail (Scheres, 2020). Nevertheless, reliable initial model generation remains the biggest hurdle in many amyloid structure determination projects.

The initial model generation consists of a 2D reconstruction of the XY cross-section of the filaments. Therefore, the results are best assessed by visual inspection of 2D images, rather than the 3D model. The output files `rec.spi` (the 2D reconstruction), `before_reproject.spi` (the summed 2D class averages along the cross-over) and `after_reproject.spi` (the projected 2D reconstruction along the cross-over) are over-written at every iteration and can be monitored with a display program that re-reads the images every time they change on disk (we use Xmipp-2.4 (Scheres *et al.*, 2008)). The algorithm should converge, with few changes to these images in the last iteration. The rec.spi image of good models typically has higher contrast (white signal against a black background) than that of suboptimal models. Good models also show continuous main-chain density, possibly even with density for bulky side chains. Multiple disconnected densities and streaks of density that extend into the solvent area are typical of suboptimal solutions. The `before_reproject.spi` image should have 2D class average images along the entire cross-over, without large discontinuities between them, and the `after_reproject.spi` image should resemble the `before_reproject.spi` image.

2D class averages displaying filaments that are not centred in the Y-direction or that are not oriented horizontally can be aligned during the 2D reconstruction process using the arguments `--search_shift, --search_angle` and `-- step_angle`. In difficult cases, pre-alignment of the images (in Relion or other software) may give better results (Scheres, 2020). If the `rec.spi` image appears symmetrical, rotational symmetry can be imposed (using the `--sym` argument) to aid convergence. Using a mask on the 2D reconstruction (`-- mask_diameter`) and limiting the resolution of the 2D reconstruction (`--maxres`) speed up the calculations and may facilitate convergence. The program has also been parallelised (`--j`), with multiple threads each aligning subsets of the 2D class averages. Quickly performing multiple runs and interactively monitoring their results facilitates parameter tuning. The most important variables to tune are the `-- crossover_distance` parameter (because it may be difficult to estimate the cross-over distance from the micrographs or the 2D class averages) and the selection of which 2D class averages to use (because it may be difficult to recognise distinct filament types).

#### 3D refinement

Next, the initial model is used for 3D auto-refinement. The initial low-pass filter applied to the initial model varies with the quality of the model, but is typically around 10 Å. The initial model is not on the same grey scale as the particles extracted for the refinement. At this stage, a helical rise of 4.75 Å (or the adjusted value if the pixel size calibration was off, see the Pitfalls section below) is used, together with a helical twist that is calculated as (4.75 Å × 180°) / *d*, with *d* being the cross-over distance (in Å) as measured in the micrographs. Typically, no point group symmetry is applied during the first refinement. Because the resolution of the model does not yet extend beyond 4.75 Å, helical twist and rise parameters are also not refined at this stage.

Refinement should result in a substantial gain in resolution over the initial model and, more importantly, result in amyloid-like features in the map, including separated β-strands and connected main-chain density with convincing side chains. If this is not the case, a better initial model may be necessary.

If additional symmetry is present in the map, the symmetry operators need to be determined. For example, if two symmetrical protofilaments are visible, the individual molecules could be related by C2 point group symmetry, or by pseudo-2_1_ helical screw symmetry. If these symmetries cannot be distinguished from the reconstruction, subsequent refinements with either of these options should be explored. The correct symmetry will lead to an increase in resolution, whereas the incorrect symmetry will prevent the map from acquiring good separation of the β-strands.

Once refinement with good β-strand separation is achieved, the helical twist and rise may be optimised. For this purpose, an initial reference with good β-strand separation is used for another 3D auto-refinement job with an initial low-pass filter close to 5 Å. To avoid overfitting, one needs to provide one of the two half-maps from a previous refinement as the initial reference. How initial models are dealt with in 3D auto-refinement has changed in Relion-4.0. If the filename contains the substring `half1` or `half2`, then both half-maps are read and set as the initial models for the two separate half-map refinements. In previous versions of Relion, the same initial model was always used for both halves. This could lead to severe overfitting, as explained in the Pitfalls section below.

In rare cases, typically with relatively noisy data, the Sidesplitter program (Ramlaul *et al.*, 2020), which is invoked through the `--external_reconstruct` argument, leads to better reconstructions than the default auto-refinement algorithm.

#### 3D classification

The separation of particle images into structurally homogeneous subsets does not work as well for amyloids as it does for single-particle analysis of globular proteins. Nevertheless, 3D classification is useful for the separation of filaments types that are relatively similar to each other, in particular when used without further alignment of the individual particle images. Varying the regularisation parameter (T=4-100) may help. One particularly useful application of 3D classification is the identification of suboptimal particles, which tend to separate from the good particles into different classes. Again, including fewer, better particles often yields better reconstructions than using more, suboptimal ones.

#### Particle polishing and CTF refinement

Beam-induced motion correction by particle polishing is typically done earlier in the structure determination process of amyloids than it is for globular proteins. The rationale behind this is that beam-induced motions can often be detected even when the reference map does not yet show all the expected features of an amyloid. If β-strand separation in early refinements is suboptimal, it is often helpful to perform polishing prior to 3D classification. Early polishing results in an early increase in the signal-to-noise ratio of the particles, which facilitates subsequent refinements. If deemed necessary, the polishing can be repeated once a better map is available.

CTF refinement, in particular optimisation of the defoci of individual particles and astigmatism for micrographs may lead to further increases in resolution. Optimisation of higher-order optical aberrations may be affected by the absence of signal at spatial frequencies in between the helical layer lines and may require better data than for globular proteins. If attempted, visual inspection of the colourful phase difference images in the log files is recommended. If a large gain in resolution is achieved in the 3D refinement after CTF refinement, executing a second CTF refinement, in particular optimising the defoci of individual particles, may further improve resolution.

#### Post-processing

Resolution estimation based on Fourier Shell Correlation (FSC) between the two independently refined half-maps and sharpening of the final map are performed in the post-processing step. A soft mask is generated using the ‘Mask creation’ jobtype, with the central Z length set to 20 or 30% of the box size. Post-processing for amyloids is run in the same automated manner as for globular proteins. However, although the resulting map will have some helical averaging applied in the Fourier domain, the real-space map will not obey helical symmetry. To impose the latter, the post-processed map is symmetrised using the `relion_helix_toolbox` program.

Because real-space symmetrisation leads to a further increase in the signal-to-noise ratio of the map, the estimated resolution from the post-processing tends to be somewhat under-estimated. This is preferred to over-estimating the resolution, which would result from convolution effects if one would attempt to measure resolution from half-maps that are symmetrised in real-space. To maximise the information content in the map, it is sometimes useful to run additional post-processing jobs, fixing the map sharpening B-factor to the value obtained from the first, automated run, but applying a higher-resolution ad-hoc low-pass filter than the resolution estimated in the automated procedure. Caution is needed when interpreting these maps, as under-filtering may lead to high noise levels. The reported resolution of the map should be the one estimated by the automated procedure.

### Pitfalls

#### Getting stuck in local minima

This probably continues to be the largest pitfall of amyloid structure determination. How to recognise and circumvent getting stuck in local minima has been described in detail previously (Scheres, 2020). The possibility of ending up in local minima of refinement means that the user needs to remain highly critical of unexpected features in the map. Although this is equally true for single-particle analysis of globular proteins, artefacts with important implications for their interpretation are much more common with amyloid reconstructions. Continuing developments in the field, like better detectors or more robust optimisation techniques, will hopefully ameliorate this situation in the future.

#### Incorrectly calibrated pixel size

In many electron microscopes, the calibrated pixel size deviates from the correct value by several percent. Incorrectly calibrated pixel sizes will lead to deviations from the expected helical rise of 4.75 Å. Such deviations can be detected from 2D class averages with sufficient resolution to separate the β-strands (looking for a peak in the spectral signal-to-noise of those images in the `model.star` file), or from the optimised helical rise in 3D auto-refinements. If an incorrectly calibrated pixel size is suspected, processing may be continued with the incorrect pixel size, but the helical rise in subsequent steps will need to be adjusted accordingly. The correct pixel size can be provided at the end of processing, through the ‘Post-processing’ jobtype, to generate a final map at the correct scale. Only for cases where the resolution extends substantially beyond 2 Å would this procedure be suboptimal, as at those resolutions higher-order effects start to become significant. Another, computationally more expensive option would be to restart processing with the correct pixel size from the beginning (although picking results could be re-used).

#### Estimating cross-over distances with higher-order symmetries

For filaments with higher than 2-fold additional symmetry, it may be difficult to estimate the cross-over distance from alternating patterns of the width of the filaments in the micrographs. An example of this is shown for the second data set described in the Results section. Therefore, when calculating the initial model in the `relion_helix_inimodel2d` program, it may be useful to attempt a wider range of cross-over distances than suggested by the micrographs.

#### Overfitting

In overfitting, noise artefacts in the reference map lead to systematic errors in the particle orientations. The artefacts are then enhanced in subsequent reconstructions with the incorrect orientations. Artefacts are most likely to appear at high spatial frequencies, where signal-to-noise ratios are low. In 3D auto-refinement, the iterative deterioration of the reconstruction is prevented by refining two maps independently against two halves of the data and low-pass filtering both half-maps at every iteration based on the FSC between them (Scheres and Chen, 2012).

Previous versions of Relion used the provided initial model as the reference for both half-sets and relied on the user to choose a suitably low-resolution initial low-pass filter to prevent overfitting. However, as described above, it is often necessary to provide relatively high-resolution initial models for successful optimisation of the helical twist and rise parameters. By providing a single, high-resolution initial model, the benefits of refining two half-sets are diminished. To address this problem, Relion-4.0 sets pairs of independently refined half-maps as initial models for the two half-sets, provided the input filename contains a `half1` or `half2` substring. Because procedures in the previous versions of Relion are amenable to accumulating high-resolution artefacts in the maps, users are urged to upgrade to Relion-4.0 and use only half-map references going forward.

A related problem exists with performing 3D classification with a single class as an alternative to 3D auto-refinement (Guenther *et al.*, 2018). In 3D classification, iterative overfitting is not prevented by separation of the data set into two halves. Instead, resolution is estimated from the power spectrum of the map itself, with higher values of the regularisation parameter T leading to higher resolution estimates. Thereby, high-resolution artefacts in the map may lead to inflated resolution estimates and the further accumulation of noise. We therefore strongly advice against this use of 3D classification. If 3D auto-refinement does not give the expected resolution of the final map, we note that the 3D auto-refinement job also responds to the regularisation parameter (through providing `--tau_fudge` as an additional argument). Using values higher than 1 will lead to higher resolution estimates during refinement, which in rare cases may improve convergence. However, as both half-maps are still refined independently, an estimate of the true resolution may still be obtained by post-processing.

#### Z-shifted half-maps

Independent refinement of two halves of the data may lead to a shift between the two maps in the (Z-) direction of the helical axis. This will lower the FSC between the two maps and thus hamper convergence onto a high-resolution solution. When the two half-maps are provided again as initial models for subsequent refinements (as described above), it will be difficult to escape from this situation. In such cases, one may align the two half-maps with respect to each other and replace one of the original half-maps with the aligned version before performing the next refinement. We use UCSF Chimera (Pettersen *et al.*, 2004) or ChimeraX (Pettersen *et al.*, 2021) for this alignment.

#### Handedness

Because cryo-EM reconstruction does not provide information on the absolute hand, the final map may need to be inverted. Most amyloid filaments solved to date have a left-handed twist, but filaments with right-handed twists have also been observed, including for filaments extracted from diseased tissue (Kollmer *et al.*, 2019; Arseni *et al.*, 2021). At resolutions beyond 2.9 Å, the handedness may be determined directly from the map through densities for the carbonyl oxygens of the main chain. For maps at lower resolutions, handedness may be inferred from the conformation of parts of the structure that have been observed previously, or from additional experiments, like atomic force microscopy or rotary shadowing electron microscopy. If this is not possible, one may also build a model in maps of both hands and compare the corresponding FSCs between the models and the maps, but it may be safer to explicitly state that the handedness remains unclear.

## Results

### Automated filament picking for in vitro assembled tau filaments

We used the modified version of Topaz for automated picking of 38 data sets in our work on *in vitro* assembly of recombinant tau (Lövestam *et al.*, 2021). For training the Topaz neural network, we manually picked 305 filaments from 44 micrographs of a data set on a tau construct spanning residues 305-379. From the resulting start-end coordinate pairs 18,094 individual particle images were extracted, using an inter-particle distance of three β-rungs, i.e. 14.2 Å. The coordinates of the individual particles were used for training the Topaz neural network, specifying a particle diameter of 120 Å and 300 expected particles per micrograph. Example micrographs with manually picked filaments used for training and automatically picked filaments by the modified Topaz approach are shown in **Figure 1**. The resulting neural network model was not only used to auto-pick the remainder of the micrographs of that same data set, but also for 37 other data sets, leading to reconstructions with resolutions ranging from 3.5 to 1.9 Å. Datasets where Topaz failed typically had large numbers of non-twisting filaments or low contrast due to thick ice or imaging too close to focus. Although non-twisting filaments may be separated by 2D classification, picking the twisted filaments might require lower thresholds, which would result in picking empty regions or the rim of the hole of the grid.

### Processing of data set EMPIAR-10940

We consider this dataset to be relatively easy, due to a readily discernible cross-over distance, and the presence of a single filament type with additional pseudo-2_1_ symmetry. Micrographs were processed as described above; auto-picked using the Topaz module with a threshold of −6 (job007 in the EMPIAR entry); and extracted using a box size of 768 pixels, downscaled to 128 pixels, with a downscaled pixel size of 4.94 Å (job009). 2D classification indicated the presence of a single filament type (job010). Four images from the 2D classification were selected (job011) to generate the initial model using a cross-over distance of 720 Å (**Figure 3** and the `inimodel/` directory in the EMPIAR entry). The initial model was rescaled to a box size of 384 pixels and the original pixel size of 0.824 Å using `relion_image_handler` and used for 3D refinement, with a twist and rise fixed to −1.19° and 4.75 Å, respectively (job014). The resulting map showed clear β-strand separation and the presence of two protofilaments related by pseudo-2_1_ helical symmetry. Subsequent 3D refinements with optimisation of the twist and rise and imposed symmetry (job017, job019), polishing (job022) and CTF refinements (job025, job026, job027) further increased the resolution. The resolution of the final map was calculated by applying a soft mask consisting of 20% of the box size (job015) and estimated to be 2.2 Å using standard post-processing (job033).

**Figure 3:**
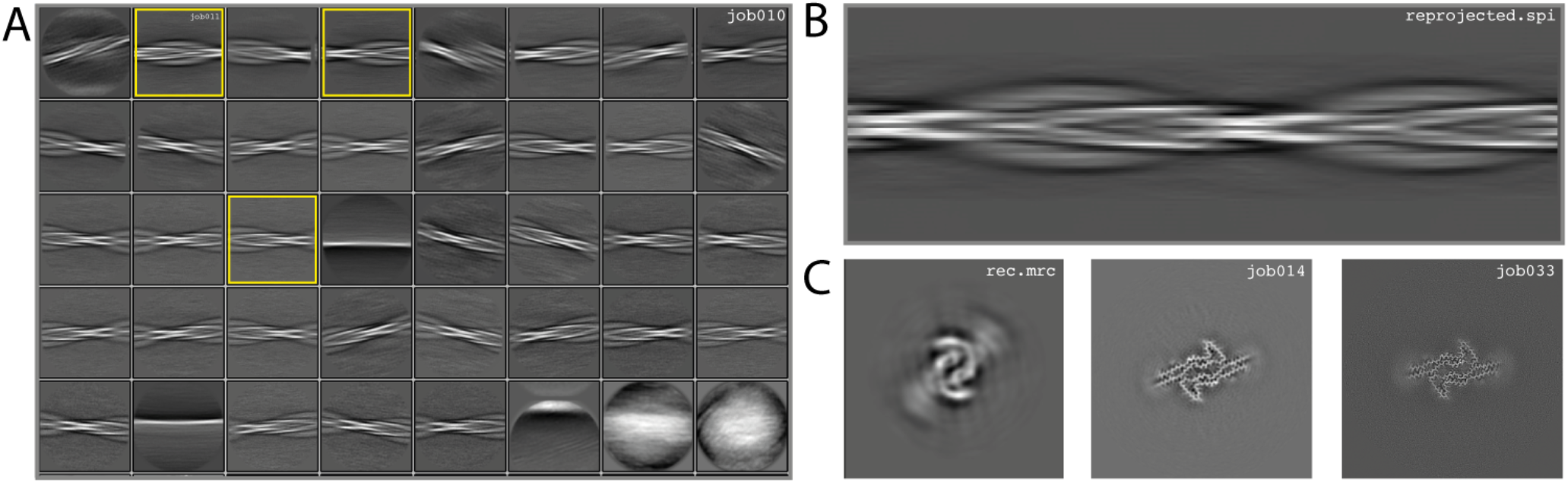
Results for data set EMPIAR-10940. **A.** 2D class averages from job010, with selected 2D class averages from job011 highlighted in yellow. The selected 2D class averages were used for initial model generation. **B.** Reconstructed cross-over from the initial model generation program. **C.** Reconstructed xy-slice from the initial model generation program (left); xy-slice from the refined map of job014 (middle) and xy-slice from the final, postprocessed map of job033 (right.)

### Processing of data set EMPIAR-10943

We consider this dataset to be of moderate difficulty, due to the presence of two filament types. The first type consists of two protofilaments without symmetry; the second type consists of three protofilaments related by C3 symmetry. Micrographs were processed as described above, auto-picked using the Topaz module with a threshold of −7 and −5 (job008 and job033 in the EMPIAR entry) and extracted using a box size of 768 pixels, downscaled to 128 pixels with a downscaled pixel size of 4.94 Å. 2D classification (job035) indicated the presence of two types of filaments, with the majority type containing a fast-twisting morphology (type A), likely with symmetry, and the other type twisting more slowly, without indications of symmetry from the 2D class averages (type B). The two different types were selected and processed separately.

Type A: Three 2D class averages were selected (job054) to generate an initial model using a crossover distance of 700 Å (**Figure 4** and the `inimodel/` directory in the EMPIAR entry). The initial model indicated that this filament is related by three-fold symmetry. As such, an additional inimodel was generated using the previous inimodel as a reference (using the `--iniref` argument), imposing three-fold symmetry (`--sym 3`) and adjusting the crossover distance to 750 Å. The resulting initial model was rescaled to a box of 384 pixels and the original pixel size of 0.824 Å using `relion_image_handler`. This model was used for 3D refinement, without additional symmetry and with a twist and rise fixed to −1.14° and 4.75 Å, respectively (job057). The resulting map showed clear β-strand separation and the presence of three protofilaments related by C3 helical symmetry. Subsequent refinements with optimisation of the twist, rise and imposed symmetry (job057, job066), 3D classification (job072), polishing (job083) and CTF refinements (job096, job097, job098) further increased the resolution. The resolution of the final map was estimated to be 2.1 Å using standard post-processing (job101).

**Figure 4:**
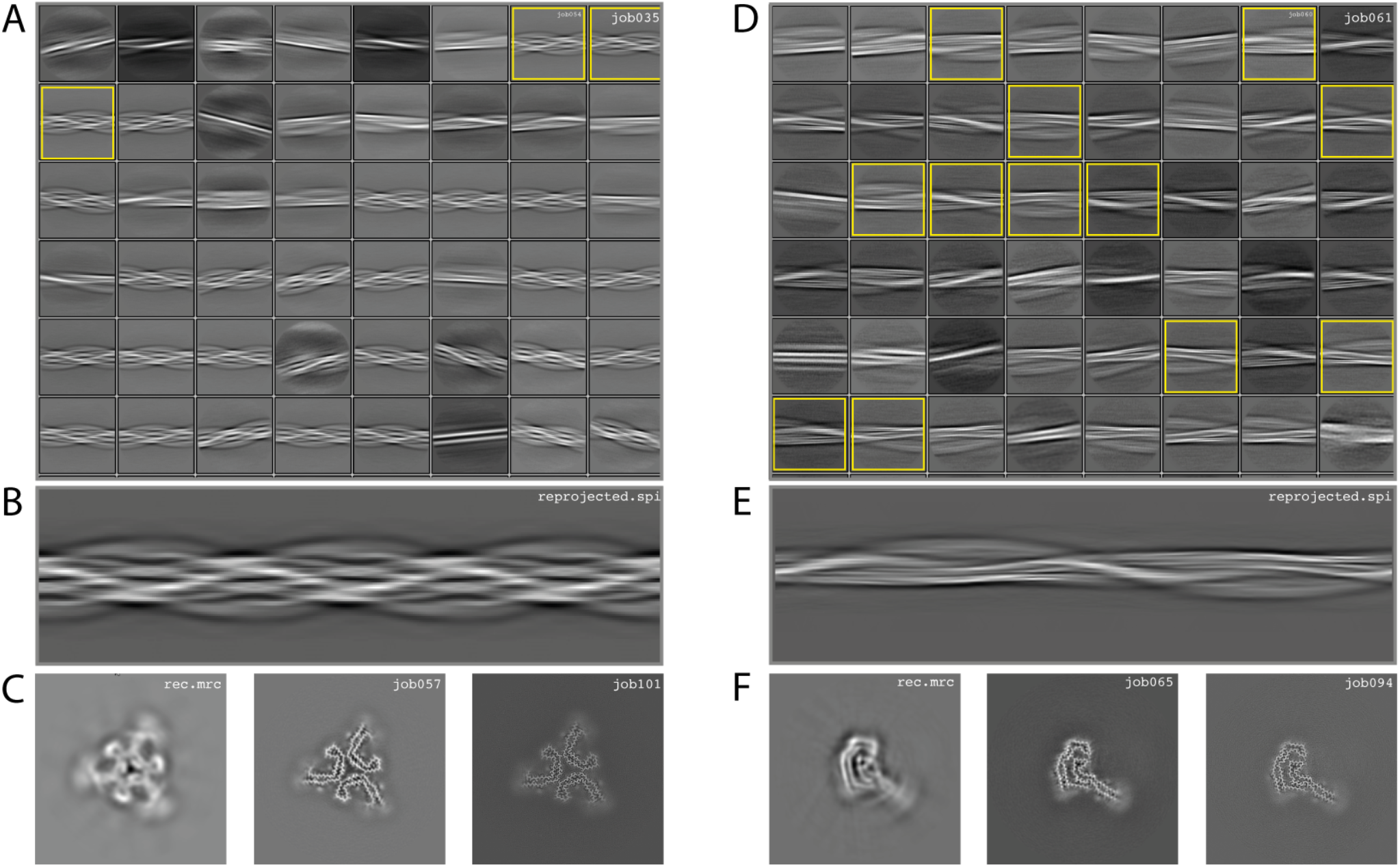
Results for EMPIAR-10943 dataset. **A.** 2D class averages from job035, with selected 2D class averages for type A filaments from job054 highlighted in yellow. The selected 2D class averages were used for initial model generation. **B.** Reconstructed cross-over for type A filaments from the initial model generation program. **C.** Reconstructed xy-slice from the initial model generation program for type A filaments (left); xy-slice from the refined map of job057 (middle) and xy-slice from the final, postprocessed map of job101 (right.) **D.** 2D class averages from job053, with selected 2D class averages for type B filaments from job060 highlighted in yellow. The selected 2D class averages were used for initial model generation. **E.** Reconstructed cross-over for type B filaments from the initial model generation program. **C.** Reconstructed xy-slice from the initial model generation program for type B filaments (left); xy-slice from the refined map of job065 (middle) and xy-slice from the final, postprocessed map of job094 (right.)

Type B: Type B filaments were selected (job051) from the 2D classification (job035), re-extracted in a box size of 512 and downscaled to 128 pixels for further 2D classification (job053). Twelve 2D class averages were selected (job060) to generate an initial model using a crossover distance of 900 Å. The initial model was rescaled to a box of 384 pixels and the original pixel size of 0.824 Å using `relion_image_handler`. This model was used for 3D refinement without additional symmetry, with a twist and rise fixed to −0.9° and 4.75 Å, respectively (job065). The resulting map showed clear β-strand separation and the presence of two protofilaments that were not related by symmetry. Subsequent refinements with optimisation of the twist and rise (job068, job069), and polishing were performed (job078). However, the map showed discontinuities in the main chain (job079). Performing a 3D refinement with local angular searches for the rot, tilt and psi angles were set to 5°, 7° and 10° respectively, and a range factor for local averaging of 3 (job086) improved the map. Subsequent CTF refinements (job090, job091, job092) further increased the resolution. The resolution of the final map was estimated to be 2.6 Å using standard post-processing (job094).

### Processing of data set EMPIAR-10944

We consider this dataset to be relatively difficult, due to the presence of four filament types. Although auto-picking did work for this dataset (job126, job127, job128), a low topaz threshold was required to all filament types, resulting in large numbers of false positives. Therefore, we resorted to manual picking (job005). Segments were extracted using a box size of 768 Å, downscaled to 128 pixels with a downscaled pixel size of 4.94 Å (job006). 2D classification (job008) indicated the presence of at least three types of filaments, but two of these were hard to distinguish. Two selections (job009 and job012) were used to process the filaments. The particles in job009 contained three filament types consisting of two protofilaments; the particles in job012 contained a single filament type consisting of three protofilaments. The processing of these two selections is described separately below.

Select job009: Initial model generation was unable to create a sensible initial model, due to (in hindsight) the presence of multiple filament types (**Figure 5**). However, as we were unable to discern the different filament types by 2D classification, we performed several refinements with different initial models (job011, job022, job029), for which the resulting maps appeared blurred, again indicative of a mixture of filament types. Therefore, we then performed a 3D classification with alignment and optimising for the twist and rise (job055), which resulted in a separation of three distinct filament types. The particles for each type were selected (job056, job057, job058) and refined with the maps generated by the 3D classification as initial model. Subsequent refinements for each polymorph were performed as described above, optimising for the helical twist and rise as well as symmetry for each type (job059, job060, job061), again followed by polishing (job080, job093, job118) and CTF refinement (job088, job089, job090, job101, job102, job103, job121, job122). The resolutions for the final maps were estimated to be 3.1 Å (job092), 3.2 Å (job111) and 3.8 Å (job124) using standard post-processing.

**Figure 5:**
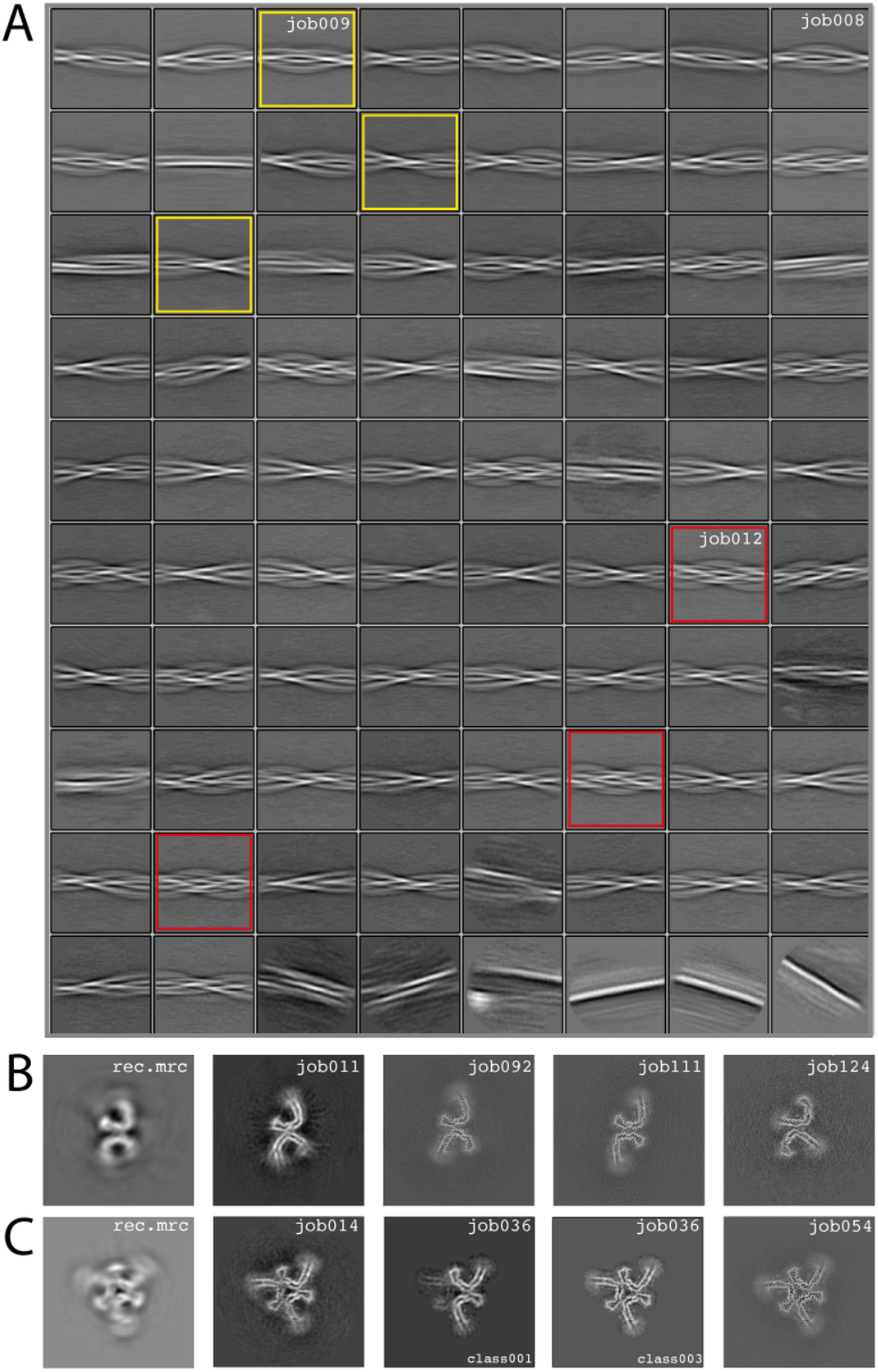
Results for EMPIAR-10944 dataset. **A.** 2D class averages from job008. 2D class averages selected in job009 are highlighted in yellow; 2D class averages selected in job012 in red. **B.** Reconstructed xy-slice from the initial model generation program for the 2D class averages selected in job009 (left); xy-slice from the refined map of job011 (second left); xy-slice of final, post-processed map of job 092 (middle), job111 (second right) and job124 (right). **C.** Reconstructed xy-slice from the initial model generation program for the 2D class averages selected in job012 (left); xy-slice from the refined map of job014 (second left); xy-slice of two classes from 3D classification job012 (middle and second right) and final, post-processed map of job054 (right).

Select job012: Initial model generation indicated this filament type consisted of three protofilaments related by three-fold symmetry, similar to the EMPIAR-10943 data set described above. This initial model was used for refinements (job014 and job020). Subsequent refinements optimising for twist and rise, and symmetry was performed (job028 and job033). The symmetry was determined by visual inspection in Chimera. Subsequent 3D classification (job036) indicated a mixture of particles containing three or two protofilaments (the latter being the same ones as selected in job092). Only particles containing three protofilaments were selected (job040) and further refined (job041). Polishing (job044) and CTF-refinements (job050, job051, job052) further improved the resolution. The resolution of the final map was estimated to be 3.2 Å using standard post-processing (job054).

## Conclusion

Amyloid structure determination is often more difficult than single-particle analysis of globular proteins. Complicated energy landscapes result in refinements getting stuck in incorrect solutions and detecting and separating multiple filament types in a data set is not straightforward. It is therefore difficult to provide a single, fail-safe procedure for automated amyloid structure determination.

But amyloids have their advantages too. They are typically more sturdy than other multi-component protein complexes objects and do not fall apart during cryo-EM grid preparation; their helical symmetry ensures excellent orientational distributions, provided the filaments twist; and every 4.75 Å of amyloid filament will yield at least one asymmetric unit for averaging. As a result, for many samples, and in particular those of recombinant protein, making grids is relatively easy and few hours of data collection often suffice for calculating reconstructions to sufficient resolution for atomic modelling.

In our experience, with some training and using the developments described in this paper, individual users can solve multiple amyloid structures per week. The automated filament picking approach described in this paper allows for full automation up to 2D class averaging using `relion_it.py` in Relion-4.0 (Kimanius *et al.*, 2021). Combined with detailed descriptions of three example data sets from our own work, we hope this will enable the use of amyloid structure determination as a high-throughput tool in many labs and look forward to the insights that these structures will bring.

## Author Contributions

S.L. performed cryo-EM structure determination; S.H.W.S. developed the automated filament picking approach in Topaz; both authors contributed to the development of image processing strategies in Relion and to writing this manuscript.

## Acknowledgements

We are grateful for T. Darling and J. Grimmett for help with high-performance computing. This work was supported by the U.K. Medical Research Council (MC_UP_A025_1013, to S.H.W.S.).

